# Solution structure of human STARD1 protein and its interaction with fluorescently-labeled cholesterol analogues with different position of the NBD-group

**DOI:** 10.1101/116368

**Authors:** Kristina V. Tugaeva, Yaroslav V. Faletrov, Elvin S. Allakhverdiev, Eugene G. Maksimov, Nikolai N. Sluchanko

**Author notes:** Corresponding author: Dr. Nikolai N. Sluchanko, A.N.Bach Institute of Biochemistry, Federal Research Center of Biotechnology of RAS, Moscow 119071, Russian Federation. 20NP: 20-((NBD)amino)-pregn-5-en-3-ol 22NC: 22-(N-(NBD)amino)-23,24-bisnorcholesterol 25NC: 25-[N-[(NBD)-methyl]amino]-27-norcholesterol 3NC: 3-(NBD-amino)-cholestane bis-ANS: 4,4’-bis(1-anilinonaphthalene-8-sulfonate) BSA: bovine serum albumin CD: circular dichroism EDTA: ethylenediaminetetraacetic acid EOM: ensemble optimization method FRET: Förster resonance energy transfer HPLC: high performance liquid chromatography IPTG: isopropyl-β-thiogalactoside LCAH: lipoid congenital adrenal hyperplasia MBP: Maltose-Binding Protein MD: molecular dynamics ME: β-mercaptoethanol NBD: 7-nitrobenz-2-oxa-1,3-diazol-4-yl PKA: protein kinase A PMSF: phenylmethanesulfonyl fluoride RALS: right angle light scattering RI: refractive index RMSD: root mean square deviation SAXS: small-angle X-ray scattering SDS-PAGE: sodium dodecyl sulfate polyacrylamide gel-electrophoresis SEC: size-exclusion chromatography StAR: Steroidogenic acute regulatory protein (=STARD1) TDA: triple detector array (Absorbance, RI, RALS).

## Abstract

Intracellular cholesterol transfer to mitochondria, a bottleneck of adrenal and gonadal steroidogenesis, relies on the functioning of the steroidogenic acute regulatory protein (StAR, STARD1), for which many disease-associated mutations have been described. Despite significant progress in the field, the exact mechanism of cholesterol binding and transfer by STARD1 remains debatable, and the solution conformation of STARD1 is insufficiently characterized, partially due to its poor solubility. Although cholesterol binding to STARD1 was widely studied by commercially available fluorescent NBD-analogues, the effect of the NBD group position on binding remained unexplored. Here, we analyzed in detail the hydrodynamic properties and solution conformation of STARD1 and its interaction with cholesterol-like steroids bearing 7-nitrobenz-2-oxa-1,3-diazol-4-yl (NBD) group in different position, namely 22-NBD-cholesterol (22NC), 25-NBD-cholesterol (25NC), 20-((NBDamino)-pregn-5-en-3-ol (20NP) and 3-(NBDamino)-cholestane (3NC). The small-angle X-ray scattering (SAXS)-based modeling and docking simulations show that, apart from movements of the flexible Ω1-loop, STARD1 unlikely undergoes significant structural rearrangements proposed earlier as a gating mechanism for cholesterol binding. While being able to stoichiometrically bind 22NC and 20NP with high fluorescence yield and quantitative exhaustion of fluorescence of some protein tryptophans, STARD1 binds 25NC and 3NC with much lower affinity and poor fluorescence yield. In contrast to 3NC, binding of 20NP leads to STARD1 stabilization and increases the NBD fluorescence lifetime. Remarkably, in terms of fluorescence response, 20NP outperforms commonly used 22NC and is recommended for future studies. Our study benefits from state-of-the-art techniques and revisits the results of the STARD1 research over the last 20 years, revealing important novel information.

## 1 Introduction

Intracellular cholesterol transfer to the mitochondrial cytochrome P450 side chain cleavage enzyme (P450scc) is considered a bottleneck of the steroidogenesis in adrenals and gonads [1-4] and determines the rate of production of pregnenolone, a single source for various types of steroid hormones. This crucial rate-limiting step relies on functioning of the cholesterol binding steroidogenic acute regulatory protein (StAR, STARD1) [2] for which many different mutations leading to severe diseases, such as lipoid congenital adrenal hyperplasia (LCAH) [1, 5, 6], have been described. In these situations, the excessive amounts of cholesterol are accumulated in the cytoplasm of steroidogenic cells that prevents production of sufficient amounts of steroids and disturbs normal cellular metabolism [1].

Human STARD1 is synthesized as a 37 kDa protein (285 residues) and contains a typical N-terminal leader sequence targeting it to mitochondria, although the 30 kDa protein devoid of this N-terminal peptide displays almost equal ability to promote steroidogenesis in COS-1 cells [7], that led to a suggestion that mitochondrial targeting either is needed to terminate STARD1 functioning or serves for some completely separate purpose [7]. The N-terminal processing of STARD1 [8], associated with the protein mitochondrial import and cholesterol transfer, seems to be regulated by cAMP levels [9] and STARD1 phosphorylation [10], and processing rates may be different in different cell types [11]. This underlines the importance of a particular cell model to be used for studies of these processes in the context of steroidogenesis [11].

The processed 30-kDa STARD1 protein comprises the so-called StAR-related lipid transfer (START) α/β structured domain (~210 amino acids), found in 15 members of the START family (STARD1-15), which forms the hydrophobic binding pocket with a size suitable for accommodation of steroids and other lipids [12]. While some START members were reported to have an affinity for different lipids like bile acids [13], ceramides and phospholipids [14, 15], steroid hormones [12], cholesterol appears the only currently known natural ligand of STARD1. Despite the crystal structure of STARD1 [16] (along with that of several other START members) has recently been solved, the structural information on cholesterol binding to STARD1 is basically confined to several, sometimes contradictory, *in silico* models lacking sufficient experimental verification. According to some predictions, the salt bridge between Glu169 and Arg188 is involved in direct cholesterol binding [17, 18] and both residues are associated with LCAH mutations [19, 20]. Mutation of many other positions in STARD1 is known to affect its functioning associated with disease conditions [5, 6, 20], albeit not always with a clear mechanistic or structural rationale. Intriguingly, despite cholesterol was used as an additive for STARD1 crystallization [16], the final structure was solved without any density for cholesterol in the putative steroid binding pocket, leaving open the question of cholesterol binding mode and orientation. Specificity of STARD1 to cholesterol analogues and other ligands is also poorly understood but these studies could be very informative for creation of artificial toxic compounds for targeting mitochondria of steroidogenic cells [21, 22].

The precise mechanism of STARD1’s action during the acute phase of steroidogenesis remains debatable, and it has long been believed that STARD1 acts at the outer mitochondrial membrane (OMM) [23, 24] and binds and releases cholesterol in a pH-dependent manner [25, 26]. This indicated that the STARD1 structure is very conformationally labile and undergoes significant rearrangements in the course of cholesterol binding and transfer, and the active protein conformation is a molten globule [17, 26]. MD simulations and studies on engineered STARD1 mutants suggested that cholesterol binding involves unfolding of the C-terminal α4 helix of STARD1 which not only endows STARD1 with the ability to bind to OMM [27], but also serves as a lid or gate for the steroid binding pocket [17, 25, 28-30]. Alternative hypothesis considered the so-called “clam-shell” like mechanism implying even more significant movements of the START domain halves necessary for the opening of the lipid binding cavity [31]. Finally, flexibility of the omega 1 (Ω1) loop was suggested to be sufficient for the lipid exchange [31, 32], but all these hypotheses required further experimental validation.

Since cholesterol is an optically inactive, poorly soluble substance, investigators used its radiolabeled [19] or fluorescently labeled [19, 33] analogues, however, radiolabeled cholesterol presents some risks and has limited availability. Fluorescent analogues of cholesterol, including commercially available 22-(**22NC**) and 25-NBD-labeled cholesterols (**25NC**) [34, 35], have been successfully used in studies of lipids trafficking and in investigations of different enzymes [35-37]. Besides obvious advantages of these reporting ligands, the main disadvantage is the presence of a relatively bulky fluorescent group which can have unpredictable effects on binding and other studied properties [34]. Although it was shown that **22NC** binds to STARD1 and its mutants similar to [^14^C] cholesterol [19], the effect of the NBD group position on the binding properties of STARD1 remained insufficiently investigated.

To fill this gap, we analyzed in detail interaction of STARD1 and cholesterol analogues with different position of the NBD group using various steady-state and time-resolved fluorescence spectroscopy approaches. To avoid previously reported problems with the STARD1 sample aggregation [20, 33, 38] and inconsistency in protein characteristics, we applied our original purification scheme [38] and obtained soluble, homogenous and monodisperse STARD1. Additionally, we for the first time comprehensively characterized its hydrodynamic properties and solution structure by small-angle X-ray scattering (SAXS) and tested previously proposed hypotheses related to the cholesterol binding mechanism.

## 2. Results and discussion

### 2.1. Purification and initial characterization of stable and soluble STARD1 species

Previously, problems with solubility of the bacterially expressed recombinant STARD1 were repeatedly reported, and the protein obtained according to traditional approaches tended to aggregate [20, 33, 38]. This arguably prevented accurate hydrodynamic characterization of STARD1 and analysis of its structure in solution. To achieve the goal, the recently described protocol, based on STARD1 fusion with a cleavable MBP-tag and providing remarkably pure, soluble, and functional protein [38], was therefore utilized. Here, the same procedure yielded milligram quantities of two STARD1 mutants, namely S195A and S195E, with modification of the residue known to be phosphorylated by protein kinase A (PKA) *in vivo* and *in vitro* [10, 39, 40]. Likewise, the phosphorylated wild-type STARD1 was obtained by protein co-expression with PKA in bacteria.

The proteins showed identical migration on SDS-PAGE (Fig. S1A), but displayed a significant shift on native-PAGE in the presence of urea (Fig. S1B), most likely due to charge differences resulting from Ser195 modifications: the S195A mutant had the lowest mobility, whereas the S195E mutant was significantly downward-shifted. The STARD1^WT^ co-expressed with PKA showed the highest electrophoretic mobility, suggesting either that the S195E substitution only partially imitates the phosphorylation state of Ser195 or the presence of additional phosphorylation sites. The latter possibility was excluded by tandem mass-spectrometry results being able to confirm only the tryptic peptide(s) ^192^RRGpSTCVLAGMATDFGNMPEQK (2449 Da with and 2293 Da without Arg192) with singly phosphorylated Ser195. Since the extent of STARD1 phosphorylation varied from batch to batch, to avoid use of heterogenous samples, we further worked with the S195A and S195E mutant proteins.

According to the CD spectra (Fig. S1C), the introduced mutations did not cause significant changes in the secondary structure of the protein, in agreement with published data [19]. Since pronounced problems with solubility were encountered in the case of the Ala mutant, whereas the Glu mutant was much more stable and could be concentrated up to 5-10 mg/ml, stored at 4 °C during several weeks, and tolerated freezing/thawing, for the following study we chose to use STARD1^S195E^ mutant. In agreement with [19], the two mutants showed similar ability to bind NBD-cholesterol (data not shown).

### 2.2. Hydrodynamic properties of STARD1

Structure of STARD6, a member of the START protein family composed of exclusively STAR domain, has been determined very recently by solution NMR (PDB entry 2MOU) [41]. It shows that the protein is monomeric and has the fold similar to that of STARD1 observed in its only available medium resolution crystal structure (residues 64-276) [16]. Despite STARD1 is considered to be highly dynamic and conformationally labile protein, potential differences in its conformation between crystallized and soluble states have not been addressed so far. Moreover, to the best of our knowledge, hydrodynamic properties of STARD1 have not been analyzed, likely due to the known propensity of this protein to aggregation.

Benefitting from the novel protocol of STARD1 production [38], we interrogated the oligomeric state of our preparation of STARD1^S195E^ by performing analytical size-exclusion chromatography (SEC) at different protein concentrations. Almost independent of protein concentration, the elution profile contained a major symmetrical peak with M_W_ = 21.1 kDa and R_H_ ~20 Å (Fig. S1D). Significantly, no aggregates were detected, however, a very small peak of dimers (<3%) appeared at increased concentrations (Fig. S1D). This indicated that our STARD1^S195E^ preparation contained unprecedentedly stable and pure monomeric species, suitable for a more detailed hydrodynamic and structural analysis. Slightly lower than expected (calculated Mw = 24.8 kDa; the electrophoretic mobility corresponded to 26 kDa (Fig. S1A)), the SEC-derived apparent size of the monomeric STARD1 (~21.1 kDa) might be explained by either some interactions with the chromatographic resin or repulsive interparticle interactions.

In order to get structural insight into the STARD1^S195E^ protein structure in solution, we performed synchrotron SAXS experiment with in-line SEC separation to ensure that the SAXS data are collected from the individual monomeric STARD1 species (see Fig. 1A). Triple detector array (TDA: absorbance, refractive index (RI), and right angle light scattering (RALS)), calibrated with BSA standard, was used to follow the elution profile and assess hydrodynamic properties of STARD1^S195E^ (Fig. 1A). This experiment confirmed the absence of protein aggregates and revealed the major peak of monodisperse particles with a very flat Mw distribution that justified further scaling and averaging of all the SAXS frames throughout the peak. The resultant curve for the STARD1^S195E^ construct (residues 66-285) (Fig. 1B) was used to assess its overall hydrodynamic parameters (Table 1). Experimentally obtained R_g_ (18.1 Å) was perfectly consistent with the corresponding parameter determined for the crystal structure of STARD1 (PDB entry 3P0L: residues 64-276, completed as described in Materials and Methods) using HydroPro software (R_g_ = 18.2 Å). Of note, this experimental value is considerably lower than the theoretical one (19.9 Å), obtained previously for STARD1 at equilibrium (starting model: PDB entry 1IMG) by MD simulations [30], and suggests a more compact protein conformation than was predicted earlier. The Mw determined by using the recently introduced volume-of-correlation (Vc) parameter [42] (in the range 0-0.3 Å^-1^) was estimated as 22.5 kDa. Porod volume (48,800 Å^3^) corresponded to near-spherical protein particles of ~28.7 kDa. Dimensionless Kratky plot with the bell-shaped peak and very small rise along the *x* axis suggested a well-folded rigid/compact domain with limited flexibility (Fig. 1C).

**Fig. 1.**
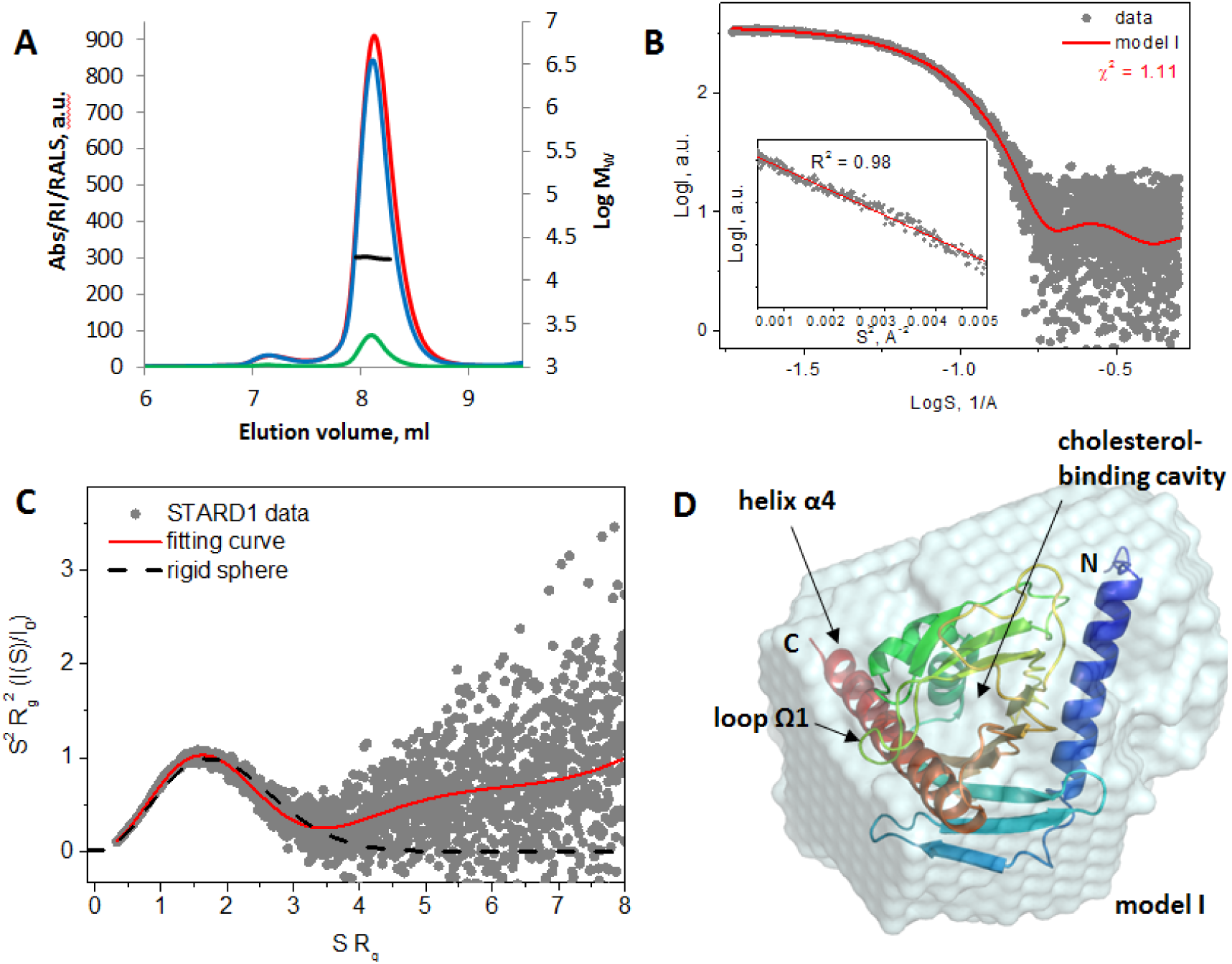
Analysis of hydrodynamics and solution structure of STARD1 ^S195E^ by SAXS coupled to SEC. **A.** SEC profile of STARD1^S195E^ followed by triple detection Malvern system (absorbance at 280 nm (Abs, blue), refractive index (RI, red), right angle light scattering (RALS, green) and synchrotron SAXS at 20 °C. Note that the flow was split for simultaneous Malvern and SAXS detection, which is reflected by a doubly reduced elution volume. Molecular weight distribution determined from RALS data and calibration by BSA run is shown by black line. **B.** The final SAXS curve produced by scaling and averaging frames corresponding to the maximum of the peak from panel A. Guinier region is shown in the inset. **C.** Dimensionless Kratky plot showing the compactness/rigidity of STARD1^S195E^ compared to the theoretical curve for a rigid sphere (black dashed line). **D.** Superposition of the average *ab initio* molecular envelope generated by DAMMIF/DAMAVER and the modified crystal structure of STARD1 (see text) providing the fit shown by red curve in panels B and C (*model 1*). The main features of the STARD1 structure are indicated. Drawn using PyMol 1.69 and Chimera 1.11. STARD1 model was superposed with the molecular envelope using “fit to map” tool in Chimera.

**Table 1.** Hydrodynamic properties of the STARD1^S195E^ mutant.

| Method used | R<sub>g</sub>, Å | R<sub>H</sub>, Å | D<sub>max</sub>, Å | Sedimentation coefficient<sup>b</sup> | Mw, kDa |
| --- | --- | --- | --- | --- | --- |
| SEC | - | 20 | - | - | 21.1 |
| HydroPro<sup>a</sup> | 18.6 | 21.9 | 63 | 2.31 S | 24.9<sup>c</sup> |
| RI<sup>d</sup> | - | - | - | - | 18.4 |
| SAXS:<br>Guinier/ P(r)/Vc | 18.1/18.4/- | - | -/59/- | - | -/28.7/22.5 |

<sup>a</sup>calculated using shell-model from atomic level on the basis of the modified crystal structure of STARD1 (PDB entry 3P0L) supplemented with missing hydrogens (residues 64-285 with extra GPGS residues at the N-terminus and S195E substitution) (see also Materials and Methods).
<sup>b</sup>calculated for 20 °C, standard conditions (viscosity and density of water solution).
<sup>c</sup>Theoretical mass of the STARD1<sup>S195E</sup> construct is 24,876 Da, SDS-PAGE derived apparent Mw is 26 kDa.
<sup>d</sup>calibrated using BSA as a standard and dn/dc = 0.185.

Together, these data provided the first hydrodynamic analysis of the monomeric STARD1^S195E^ and suggested that it can be approximated with rigid, non-aggregated and monodisperse particles, providing a prerequisite for structural modeling.

### 2.3. Model of the STARD1 structure in solution

The apparently closed STARD1 structure, with the buried hydrophobic cavity of a size comparable to that of cholesterol, has been puzzling researchers for years, and the ligand binding mechanism by STARD1 still remains debatable [41, 43]. Several hypotheses, based on preexistence of different open and closed conformations, were put forward, however, experimental evidence, that would clearly support one or another proposed binding model, continues to be lacking.

The available crystal structure of STARD1_64-276_ (PDB entry 3P0L, chain A), corrected to account for slight differences in utilized constructs (see Materials and Methods), was consistent with the *ab initio* molecular envelope built using DAMMIF/DAMAVER [44] and provided an excellent fit (χ^2^ = 1.11) to the experimental SAXS curve (Fig. 1B and D, *model I*), suggesting that the equilibrium state of STARD1 does not deal with substantial structural reorganization of the protein molecule, in contrast to the proposed earlier models attempting to explain the cholesterol-binding mechanism [18, 31]. For instance, dimensions of the open model that would facilitate cholesterol entry due to its “clam shell”-like structure, as in PITPα protein (=STARD10), were slightly larger than suggested by the experimental SAXS data for STARD1^S195E^ (χ^2^ = 2.14; Fig. S2, *model II*), and therefore such conformational rearrangements are unlikely for STARD1 in solution. Likewise, when we considered the previously proposed unfolding of the C-terminal helix α4 as a gating mechanism [17, 18, 28] and created 10,000 models to exhaustively sample different positions of the unfolded C-terminal segment (residues 250-284), we found that only the most compact models with unfolded C-terminal segment sticking to the rest of the protein were selected by the ensemble optimization method (EOM) as matching the SAXS data, making such a scenario also questionable, at least with the apo-STARD1 at pH 7.5 (Fig. S2, *model III*). This experimental observation is in striking contrast with the MD simulations data predicting the equilibrium state of STARD1 to be intermediate between the fully open (helix α4 detached and unfolded) and fully closed [18, 30]. Moreover, such a situation would inevitably lead to increased protein flexibility, which is not however supported by the Kratky plot (Fig. 1C).

On contrary, we found that suggested earlier movements of the loop Ω1, while obviously being sufficient for the opening of the cholesterol-binding cavity and ligand penetration [31], are consistent with our SAXS data. Indeed, manually built models with different position of the loop (Fig. 2, *model IV*) provided almost identical, excellent fits to the experimental SAXS curve (χ^2^ ranging from 1.11 to 1.12) and could also describe the data as an ensemble (EOM fit: χ^2^ = 1.11). The conformational fluctuations in *model IV* are also in line with the fact that in the crystal structures of various STARD homologues residues of the loop Ω1 possess either higher B-factors or slightly different spatial positions.

**Fig. 2.**
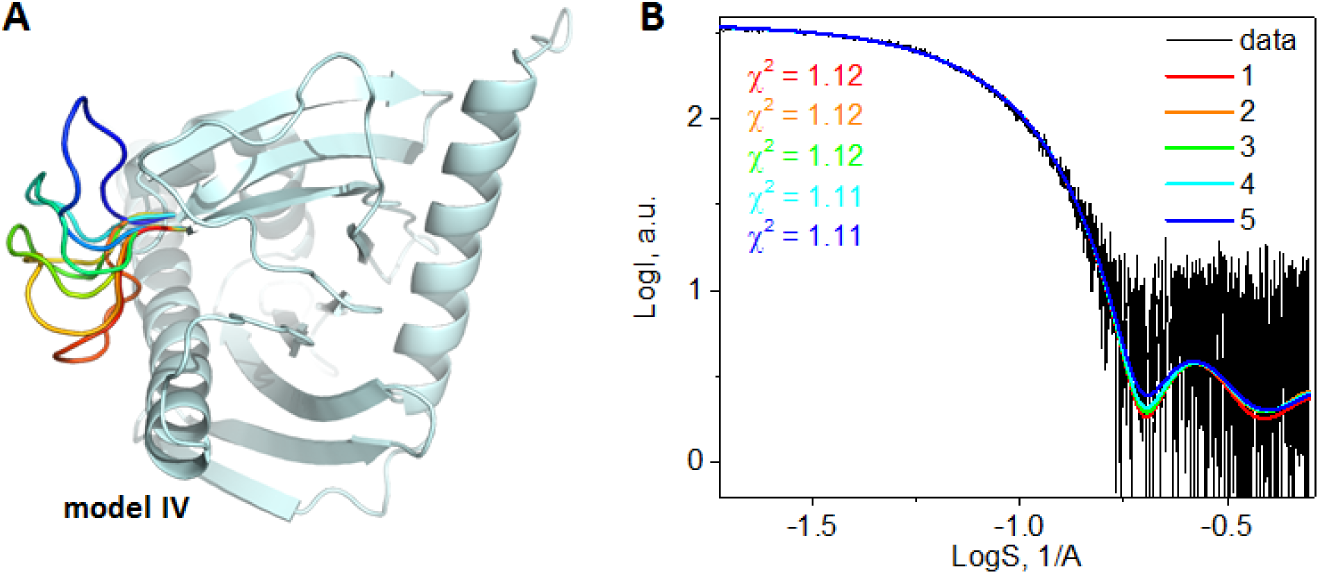
Solution structure of STARD1 considering movements of the loop Ω1 (residues 171-182) as the only conformational changes in the protein as a gating mechanism for cholesterol entry. **A.** Structural representation of model IV. Different positions of the loop are shown by color gradient from red (closed state) to blue (open state). **B.** Fits to the data with the same color coding. Drawn using PyMol 1.69, models were manually built on the basis of the 3P0L structure using Coot [45].

It is therefore unlikely that in physiological solution STARD1 experiences serious spontaneous conformational changes to facilitate cholesterol binding. Our SAXS data also supports the idea of structural similarity between STARD1 and STARD6, since the recently determined solution NMR structure of the latter, described by 20 conformationally different states with a maximal pairwise backbone RMSD of 2.55 Å (PDB entry 2MOU), provided an excellent fit to our SAXS curve (χ^2^ ranging from 1.13 to 1.16, not shown).

Notwithstanding similarity of the overall fold, the START family members are known to display significant ligand specificity, which is believed to be associated with differences in the lining of their cholesterol-binding cavities. Indeed, apart from cholesterol [46], i.e., the known ligand of STARD1, STARD3, STARD4, STARD6, but not STARD5 [43], STARD6 has recently been reported to specifically bind testosterone [41]. At the same time, STARD5 is a unique bile acids binding START family member [13]. Due to specialized linings in the START members, the ligand specificities and binding orientation cannot be simply extrapolated from one member(s) to another. To the best of our knowledge, not only physiological ligands of STARD1 besides cholesterol are unknown, but the orientation of the bound cholesterol molecule in the cavity of STARD1 is not clearly elucidated yet. In particular, it was shown *in silico* that the two main orientations of cholesterol, with the 3-OH group looking inside the cavity (mode “IN”) or toward the loop Ω1 (mode “OUT”), are equally probable [31], and therefore additional experimental information is highly required to distinguish between the two. Curiously, despite cholesterol was used as an additive to STARD1, the crystal structure of the latter was solved with no density that would support the presence of cholesterol in the binding cavity, leaving its exact binding mode unknown [16].

### 2.4. Interaction of STARD1 with fluorescent cholesterol analogues: the effect of the NBD group position

To get insight into the mode and selectivity of the cholesterol binding to STARD1, we probed the latter with a series of cholesterol analogues with the fluorescent 7-nitrobenz-2-oxa-1,3-diazol-4-yl (NBD)-group attached to either 20^th^ (**20NP**) [38, 47], 22^nd^ (**22NC**), 25^th^ (**25NC**) carbon atoms of the side chain, or to the 3-OH group (**3NC**). The structural formulae of these compounds are depicted in Fig. 3A. The NBD group is solvatochromic, i.e., reacting on the polarity of the environment by increasing the quantum yield of fluorescence and by the blue spectral shift upon translocation to the hydrophobic/masked regions. It is present in some commercially available NBD-cholesterols, e.g., 22-NBD-cholesterol (=**22NC**) [25, 48, 49]. Importantly, despite the presence of the rather bulky fluorescent group, **22NC** was shown to be essentially equivalent substrate to study STARD1 binding as the radiolabeled cholesterol [19], justifying utilization of NBD-cholesterol analogues. In line with this, we have recently reported that STARD1 is capable of binding a shorter synthetic compound, **20NP**, whose binding to STARD1 shows similar features as **22NC** [38].

**Fig. 3.**
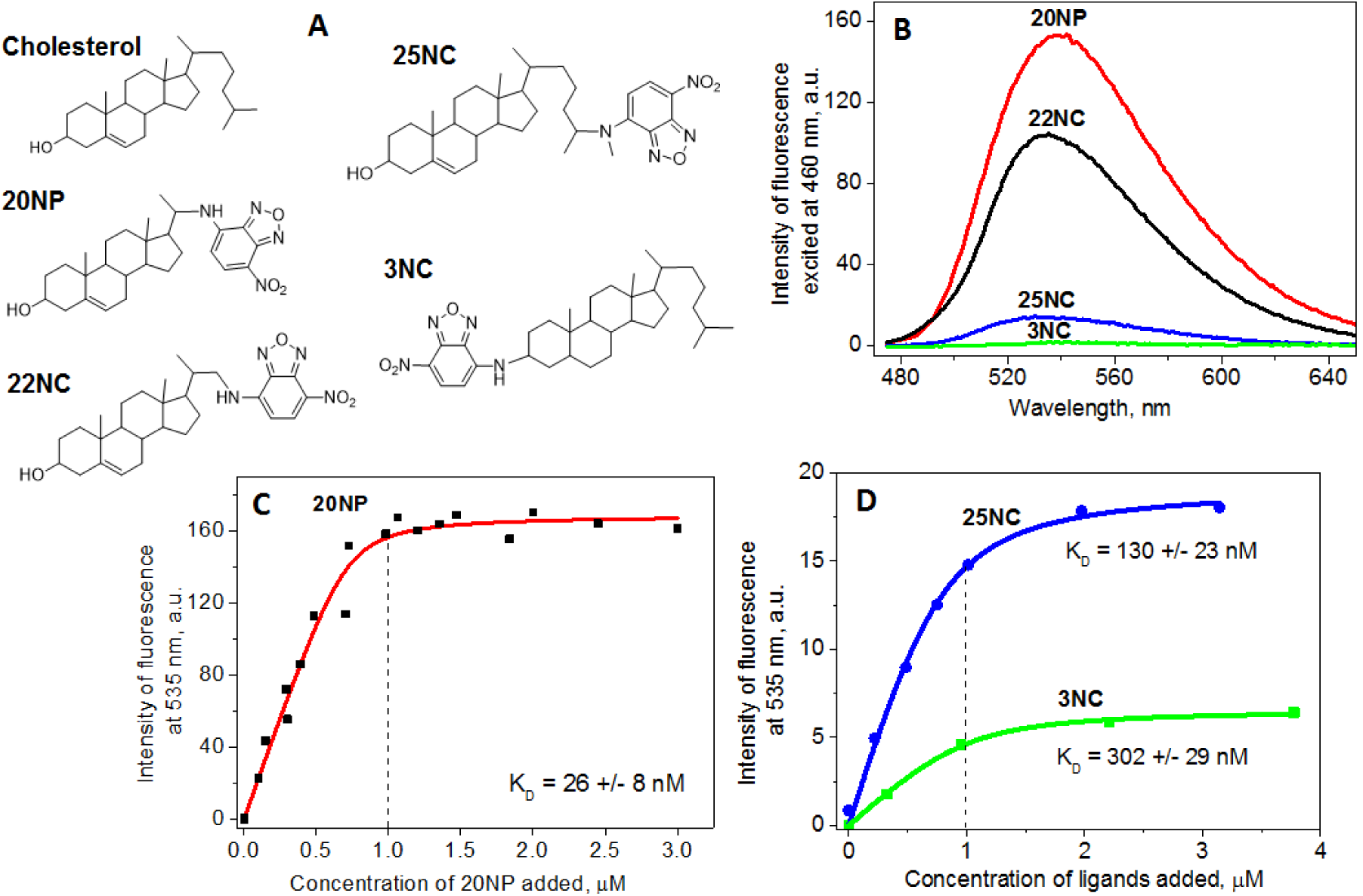
Probing STARD1^S195E^ by fluorescent analogues of cholesterol with different position of the NBD-group. **A**. Structural formulae of the ligands studied. **B**. Fluorescence response of the ligands (1.5 μM) as a result of binding to STARD1^S195E^ (1 μM). Three spectra for each ligand were collected, buffer-subtracted and averaged. **C**, **D**. Titration of 1 μM STARD1^S195E^ solution with increasing concentrations of ligands 20NP (**C**), 25NC and 3NC (**D**). Fluorescence of the NBD-labeled ligands was excited at 460 nm at slits width of 5 nm, and after pre-incubation was recorded at 535 nm at 37 °C. Binding curves were approximated with the quadratic equation [38] tolerating the difference between the total and free ligand concentrations. Vertical dashed line indicates the 1:1 ratio. The titration experiment was repeated at least five times for each ligand with the most representative results shown.

Here, we questioned whether the position of the NBD group can affect the binding and shed light on the orientation of the cholesterol core in the STARD1 cavity and probed the interaction of STARD1^S195E^ by cholesterol analogues with the NBD group located either at the side chain (**20NP**, **22NC**, **25NC**), or instead of the 3-OH group (**3NC**) which is believed to be important for cholesterol binding due to formation of the polar contacts with the residues within the cavity [50].

In the beginning, we confirmed the physical interaction of **20NP**, **22NC**, **25NC**, and **3NC** with STARD1 by analytical size-exclusion chromatography of the corresponding pre-incubated protein/ligand mixtures upon dual detection at 280 (STARD1) and 460 nm (ligands). This experiment showed that all the ligands co-eluted with the monomeric STARD1^S195E^ at 11.24 min, and no significant differences were found in the amplitudes of the peaks in case of **20NP**, **22NC**, **25NC** (Fig. S3), however, **3NC** co-eluted also with protein aggregates at ~5.50 min and, likely, stimulated formation of large associates or micelles. In this case, the amplitude at 11.24 min slightly decreased (Fig. S3), suggesting only slightly less binding of this compound to the monomeric STARD1. We would like to point out that such a hydrodynamic analysis helps reveal aggregation state and quality of the preparation and is important in the context of STAR/ligand interactions, although has not been performed previously in such studies.

We observed dramatic changes in the level of fluorescence of ligands pre-incubated with STARD1^S195E^ at a 1.5-fold ligand excess, i.e., decreasing in the row **20NP** > **22NC** >> **25NC** > **3NC**, suggesting that either parameters of their binding or those of fluorescence emission are cardinally affected by the location of the NBD group (Fig. 3B). Surprisingly, the highest intensity of fluorescence was observed in the case of **20NP**, and not commercially available **22NC** compound (Fig. 3B). Given that both ligands were reported to bind to STARD1 [38, 48, 49], this may be associated with differences in the NBD group environment and its relative position with adjacent tryptophan residues of STARD1. Noteworthy, the possible proximity of Trp241 and the NBD group in bound ligands **20NP** and **22NC**, visualized by protein-ligand docking (Fig. S4C), leads to the efficient FRET from Trp to NBD (Fig. S4 and [42]) as their spectra overlap (Fig. S4D). Given that STARD1 contains four Trp residues and assuming that they roughly equally contribute to the fluorescence intensity, the observed Trp fluorescence quenching ((F_STARDI_-F_STARDI/2ONP_)/F_STARDI_) upon binding of **20NP** indicates that one Trp could be quenched completely. We assume that the most proximal Trp241 may be fully quenched by bound **20NP** as the distance between the donor and acceptor does not exceed 12 Å, confirming specific binding of **20NP** in the cavity of STARD1. Importantly, the FRET changes upon **20NP** addition were saturable, yielding ~20 nM apparent dissociation constant (Fig. S4B, and see below). These observations are qualitatively in line with the previously reported ones [49], however, the emission of the UV fluorescence maximum of STARD1 in [49] (352 nm) is ~15 nm red-shifted compared to our results (~338 nm; Fig. S4A), and the efficiency of FRET in [49] is significantly lower (64%) than in our case (ca. 100%). This points to significant differences in protein preparations and, given the known propensity of STARD1 to unfolding and aggregation, especially for refolded STARD1 preparations [38], emphasizes the necessity of exhaustive hydrodynamic characterization of STARD1 preparation before ligand binding assays.

Although the intensity of fluorescence of **20NP** bound to STARD1 was 10 or 20 times higher than that of **25NC** or **3NC**, respectively (Fig. 3B), when STARD1^S195E^ was titrated by increasing amounts of these ligands (Fig. 3C and D), we observed binding curves with saturation at high concentrations in all three cases. These suggested rather specific interaction occurring at a ~1:1 stoichiometry (as in [19, 25, 33, 38, 51]) and allowed us to assess binding affinities. Upon approximation [38], we found that **20NP** binds to STARD1 with an apparent KD of 26 ± 8 nM (Fig. 3C), which is very similar to the earlier reported values for cholesterol (30 nM; one binding site) [51] and **22NC** (32 nM; two binding sites) [49], but is substantially lower than the apparent K_D_ for **3NC** (302 ± 29 nM) and **25NC** (130 ± 23 nM) (Fig. 3D). This is in line with the previous conclusion that all the tested ligands are able to bind to STARD1, although with significantly different affinities. The binding of **3NC** and **25NC** results in the very small increase in the NBD fluorescence intensity, implying significant quenching of the latter, presumably, due to its (partial) contact with the solvent.

In accordance, pre-incubation of STARD1^S195E^ with a 5-fold excess of **3NC** led to a decrease in the fluorescence response from **20NP** binding to STARD1^S195E^, confirming competition between the two for the ligand binding cavity (data not shown). The specificity of binding *within the cavity* was additionally confirmed in the experiment, in which STARD1 was first titrated by **20NP** until saturation, and then changes in the intensity of NBD fluorescence were followed upon addition of cholesterol stock solution. This led to a dose-dependent decrease in fluorescence intensity, indicating gradual replacement of bound **20NP** by cholesterol (Fig. S5).

Together, our data indicate that i) all tested ligands are able to bind to STARD1, ii) the most efficient reporter is **20NP** (having affinity comparable with that of cholesterol), iii) **3NC** and **25NC** are poor reporters of STARD1/cholesterol interaction, with marginal response and relatively low affinity. This may be associated with the larger sizes of **3NC** and **25NC** precluding them from being fully accommodated in the cavity. We cannot exclude that these large ligands can bind less specifically to some surface areas of STARD1 *(outside* the cavity). Curiously, by titration of pre-formed STARD1/**20NP** complexes with the unspecific hydrophobic dye bis-ANS, we could find that the ligand-bound form of STARD1 still shows pronounced surface hydrophobicity (unpublished observations), indirectly supporting the notion that hydrophobic ligands that are bigger than the cavity may protrude or bind outside of it.

We tried to validate our experimental observations by performing *in silico* docking of cholesterol, **20NP**, **25NC**, and **3NC** into STARD1 structure. As mentioned above, earlier it has been concluded that the two opposite orientations of the cholesterol molecule inside the STARD1 cavity, denoted as “IN” or “OUT” depending on where the 3-OH group is looking, are roughly equivalent thermodynamically [31]. Therefore, it is not clear, why the “IN” binding mode has become widely assumed to be the correct one, especially given that the experimental data have been desperately lacking. Our attempts to perform docking of the cholesterol molecule into the modified STARD1 (Fig. 1, *model I*) using either Autodock [52] or FlexAID [53] resulted in “C3 OUT” as the more preferential binding mode, and so was the case if **20NP** (or **22NC**) were docked (Fig. 4). On contrary, Autodock docking of **25NC** and **3NC** resulted in only rare poses out of the top 10 with ligands lying inside the cavity, in a random orientation (not shown), indicative of less favorable binding than in case of cholesterol and **20NP** (or **22NC**). Remarkably, docking using FlexAID supported our conclusions for cholesterol and **20NP**, however, resulted in **25NC** and **3NC** binding with the cholesterol core *inside* and the NBD group looking *outside* the cavity (Fig. 4), in line with the hypothesis that the low fluorescence response in their case is associated with the lower affinity of these ligands and quenching of their NBD groups as the result of partial exposure. Importantly, notwithstanding different scoring functions to assess binding poses [52, 53], the two algorithms complemented each other in predicting less preferential binding of **25NC** and **3NC**, but favorable and consistent binding of cholesterol and **20NP** in the “C3 OUT” mode.

**Fig. 4.**
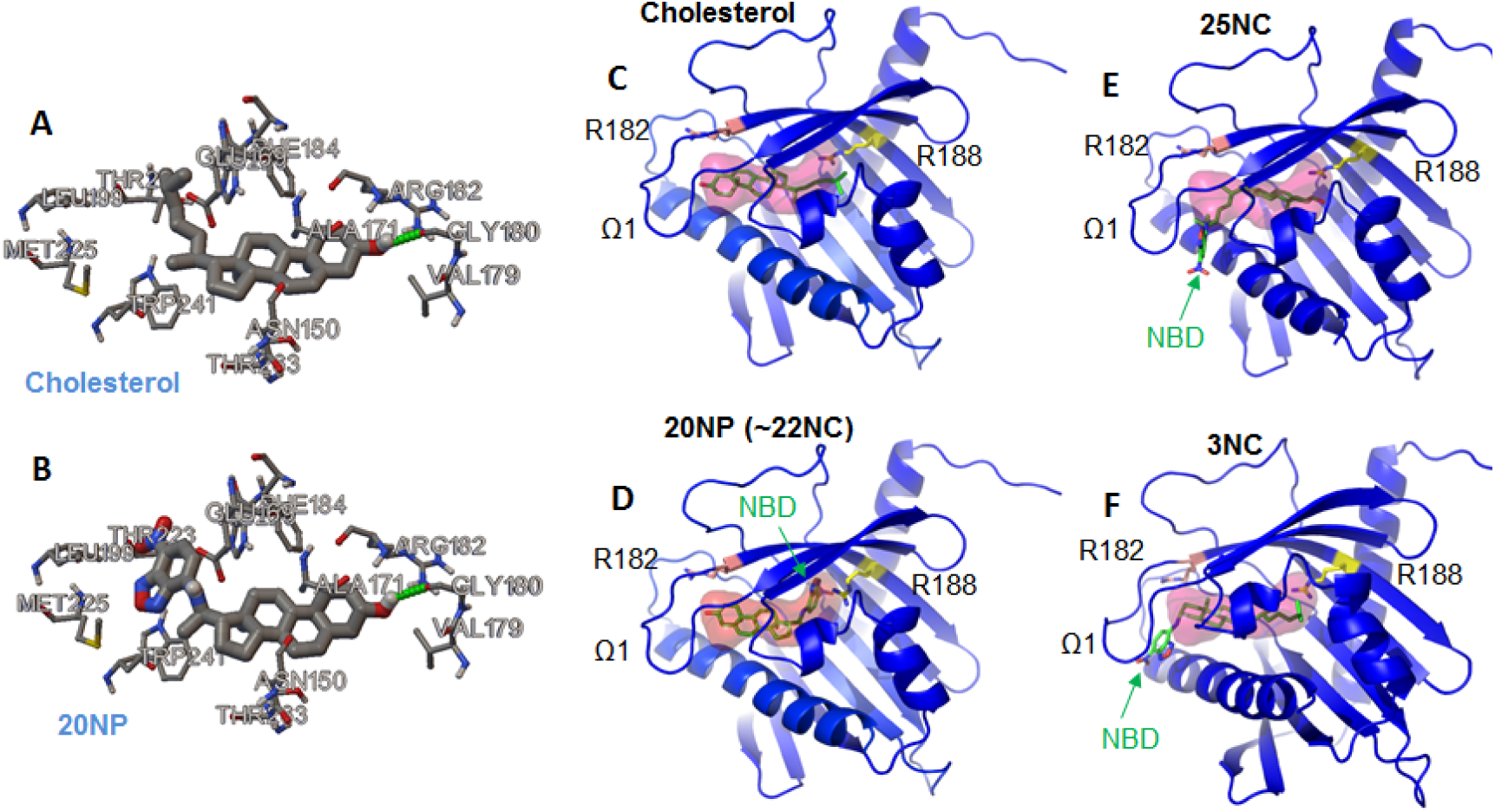
Docking of cholesterol and its fluorescent analogues using Autodock (**A, B**) and FlexAID (**C-F**). Best poses (pose “0”) for each simulation run are shown. For Autodock runs, only the results for cholesterol and 20NP (sticks and semitransparent surface, with 3-OH group marked red) docking are shown, 25NC and 3NC tended to locate outside the cavity or the results were not reproducible. For FlexAID runs, the best poses of ligands (green sticks) in the STARD1 (blue ribbon) are shown, with the binding volume (semitransparent surface), loop Ω1, NBD group (green arrow), and R182 (salmon) and R188 (yellow) residues determining the orientation “IN” (C3 carbon atom of a ligand looking at R188) or “OUT” (C3 carbon atom of a ligand looking at R182) marked. Note that in 25NC and 3NC cases, the NBD group is looking outside and is partially exposed. The docking simulations were repeated thrice and the most typical results are presented. The pictures were obtained using either Autodock tools 4.2 (A, B) or PyMol 1.69 (C-F).

It was earlier proposed that R188 forms a salt bridge with Glu169 and plays a key role in coordination of the 3-OH group of cholesterol [17, 18], whereas R182 located near the omega 1 loop may take part in regulation of its mobility [31]. Intriguingly, the published data on various STARD1 mutants with substitutions of R182 and R188 residues, often associated with pathologies like LCAH, tell in favor of the importance of these residues, but cannot help to unambiguously distinguish whether the “C3 IN” or “C3 OUT” cholesterol binding mode is the natively occurring one. Both residues are highly conserved in STARD1 orthologs annotated in the UniProt database (14 species), however, R182 residue is more conserved among human START paralogs (i.e., 15 members of the START family; e.g., STARD15 contains Lys instead of Arg in this position), indirectly implying its more universal role in ligand binding than R188. In line with this, in STARD5, the equivalent position of Arg188 is occupied by Val120 (Fig. S6), despite cholesterol and bile acids were still proposed to bind to STARD5 in the “C3 IN” orientation [16, 46].

### 2.5. Thermal stability of STARD1 and the effect of ligand binding

It is known that binding of cognate ligands can affect stability of the START family members [41, 43, 51]. We sought to investigate this phenomenon using our apparently stable STARD1 preparation obtained from the MBP fusion [38] and a combination of steady-state and time-resolved fluorescence spectroscopy.

The STARD1^S195E^ sample was pre-equilibrated at 10 °C in the absence or the presence of different ligands and then was heated up with a constant rate of 1 °C/min, and the intensity of intrinsic tryptophan fluorescence (excitation 297 nm, emission 346 nm) was measured as a function of temperature (Fig. 5 and Fig. S7). A typical thermal unfolding curve could be divided into three regions corresponding to the folded state (F), transition (T), and unfolded state (U) (Fig. S7A). Linear approximation of the first and the third regions enabled building dependencies of completeness of transition on temperature (Fig. S7B), which in turn allowed estimation of half-transition temperatures (T0.5) (Fig. 5). We found that, while T0.5 of STARD1^S195E^ was equal to 49.8 ± 0.1 °C, **20NP** increased this value by more than 3.5 degrees (T0.5 = 53.4 ± 0.1 °C), however, the presence of **3NC** did not affect the thermal stability of the protein (T0.5 = 49.5 ± 0.1 °C). Morevover, when added in an excess over **20NP**, **3NC** significantly decreased the stabilizing effect of the latter. Importantly, this supports the idea that **3NC** and **20NP** compete for the binding to STARD1 and suggests that **3NC** binding occurs with lower affinity and fails to increase the thermal stability of the protein, supporting our previous conclusions. The presence of cholesterol also increased the thermal stability of STARD1 (from 49.8 ± 0.1 °C to 50.7 ± 0.1 °C), although to somewhat lesser extent than in the case of **20NP** (T0.5 = 53.4 ± 0.1 °C).

**Fig. 5.**
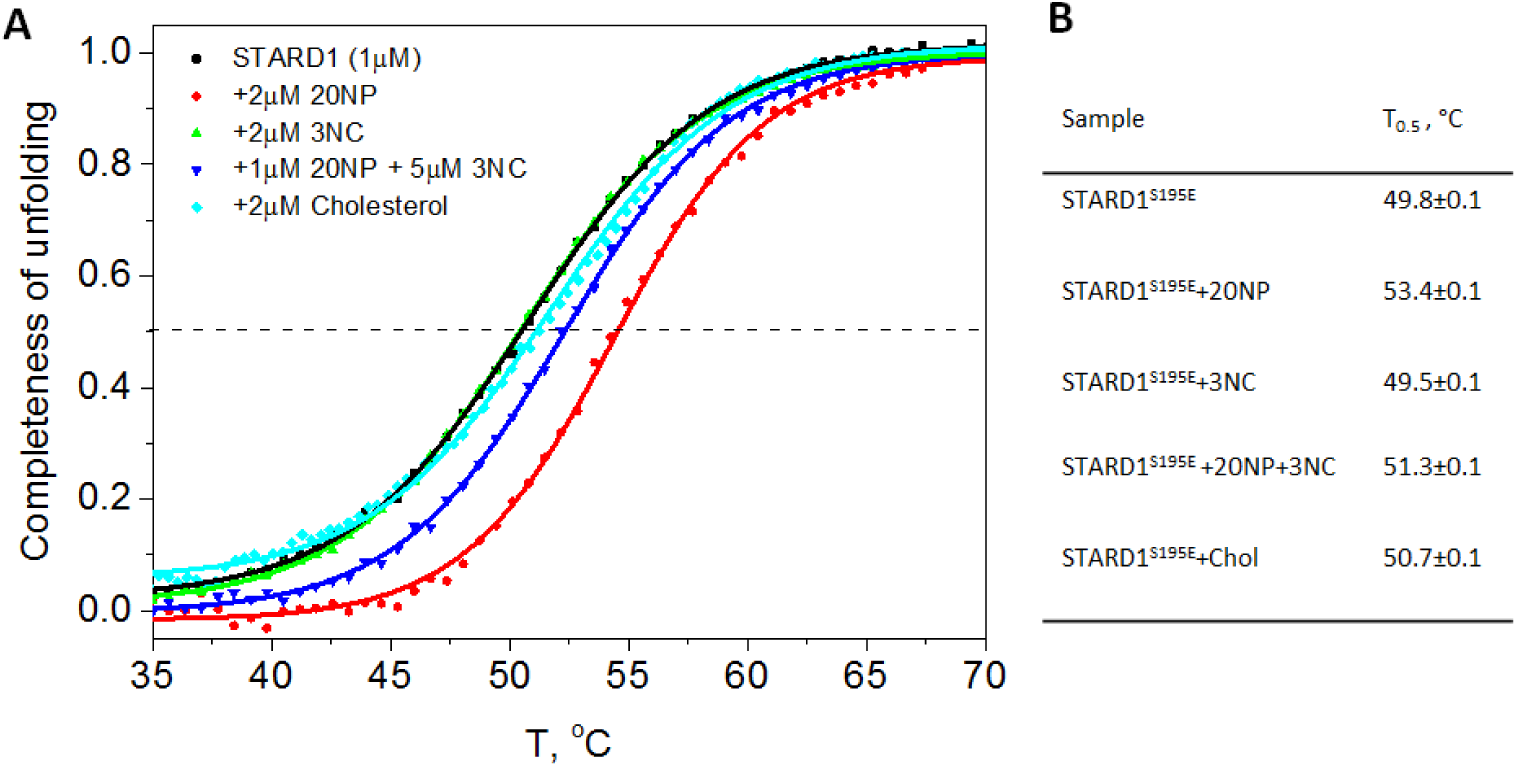
Thermal stability of STARD1^S195E^ in the absence and in the presence of its ligands followed by changes in intrinsic tryptophan fluorescence of STARD1. **A**. The temperature dependences of completeness of unfolding for the samples as indicated. The expeiment was done four times and the most typical results are presented. **B**. The T_0.5_ values for these samples determined from the panel A. See Fig. S7 for more details.

We would like to underline that the T_0_._5_ value for our STARD1 preparation (49.8 ± 0.1 °C in the apo-form) obtained using the MBP fusion [38] is significantly higher than that corresponding to STARD1 preparations obtained in earlier studies by using CD spectroscopy [51] (T0.5 = 42.3 ± 0.1 °C in the apo-form). In spite of similar observed stabilizing effect of the ligand binding in both studies, this points to potential differences in the quality of preparations and emphasizes the necessity of exhaustive preliminary characterization of the protein before binding studies.

In the absence of STARD1, fluorescence lifetimes (short τ_1_ and long τ_2_) of the solvent exposed NBD group in **20NP** (τ_1_ = 225 ps (79.7%); τ_2_ = 3435 ps (20.3%)) and **3NC** (τ_1_ = 295 ps (72.2%); τ_2_ = 2980 ps (27.8%)) were low (Fig. 6A). Upon addition of the STARD1^S195E^ excess, only the lifetimes for **20NP**/STARD1^S195E^ significantly increased (τ_1_ = 2185 ps (55%); τ_2_ = 9860 ps (45%)), those for **3NC** remained almost unchanged (τ_1_ = 235 ps (76.1%); τ_2_ = 4145 ps (23.9%)) (Fig. 6A). This suggests that the NBD group of **20NP** becomes protected upon interaction with STARD1, whereas **3NC** remains almost equally quenched in the free and bound states. This is in line with the conclusion that the latter ligand displays lower affinity to STARD1 and, upon binding, cannot be accommodated in the cavity in full (Fig. 6C), resulting in the exposed NBD group and very low fluorescence response. It may also be possible that the yield of the slow component of **3NC** fluorescence increases in presence of STARD1 partially due to its ability to form large associates and co-aggregate with the protein (Fig. S3B).

**Fig. 6.**
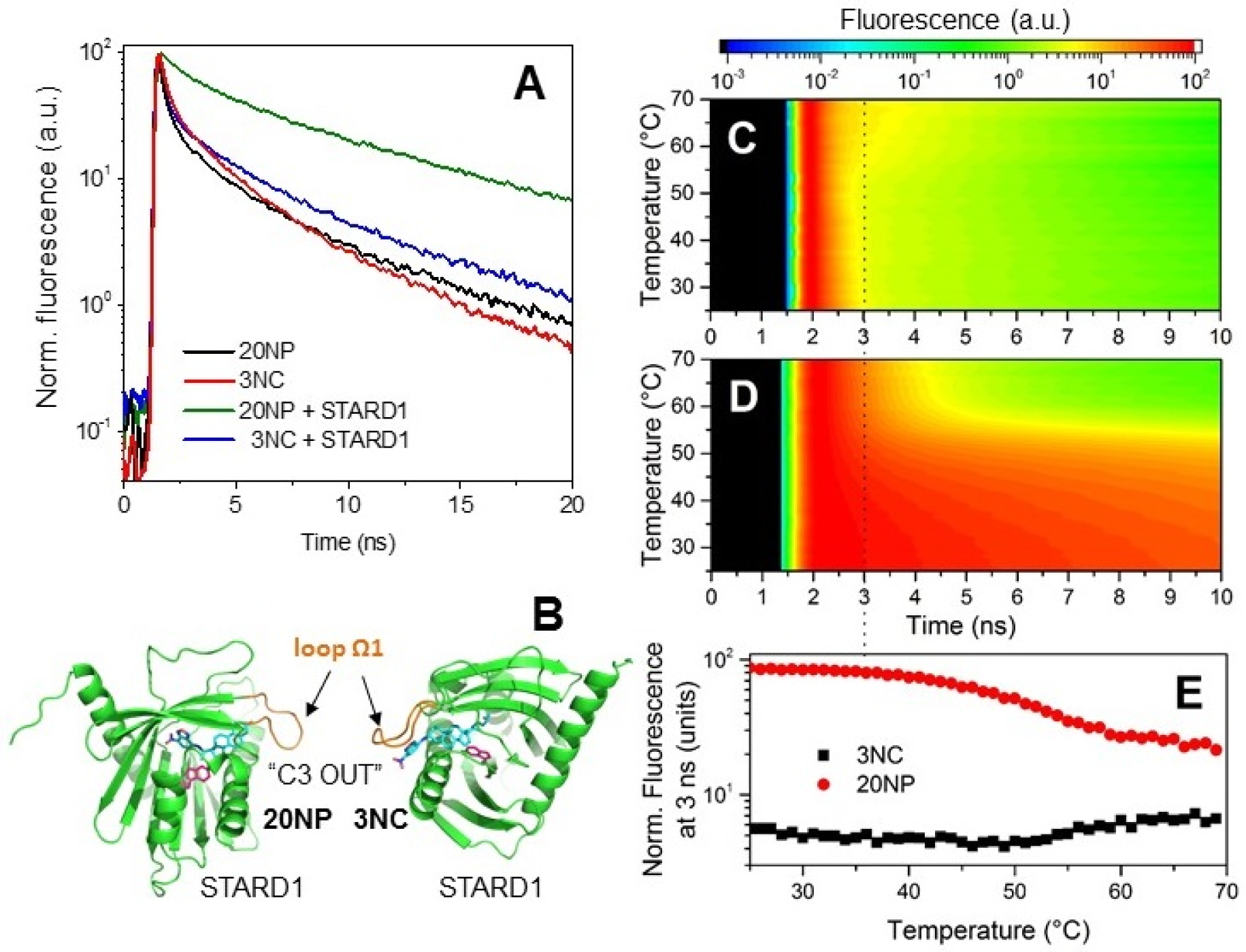
Time-resolved fluorescence spectroscopy study of the STARD1^S195E^ interaction with 20NP and 3NC ligands. **A**. The NBD fluorescence decay kinetics at 25 °C for 20NP and 3NC ligands (1.4 μM) in the absence and in the presence of STARD1^S195E^ (2 μM). **B**. Hypothetical models of the STARD1^S195E^ interaction with 20NP and 3NC as predicted by FlexAID docking. The best scoring poses, consistent with the experimental data in the study, are shown. Ligands are shown in cyan, Trp241 of STARD1 (green) is shown in magenta, loop Ω1 is orange. **C**, **D**. Color maps demonstrating the thermal stability of STARD1 in the presence of 3NC (**C**) 20NP (**D**) as studied by changes in the fluorescence decay kinetics of the NBD group during the heating of the samples from 25 to 70 °C at a constant rate of 1 °C/min. **E**. Profiles of fluorescence intensities at 3 ns (proportional to the slow component contribution, i.e. reflecting the behaviour of the protein-bound ligand) after fluorescence excitation by a picosecond laser flash, corresponding to the dashed line in (**C** and **D**). The experiment was done twice with the same results.

Our conclusions were further confirmed by the experiment, in which pre-incubated STARD1^S195E^ mixtures with either **20NP** or **3NC** were heated with a constant rate of 1 °C/min with continuous registration of the NBD fluorescence decay kinetics. This allowed to visualize changes in the lifetimes of the NBD fluorescence and additionally analyze the thermal stability of STARD1 by unfolding-induced dissociation of fluorescent ligands (Fig. 6). In agreement with Fig. 6A and above, at temperatures below 50 °C, the fluorescence lifetime was dramatically larger in the case of **20NP**, and gradually decreased upon heating. The sharp decrease in its fluorescence lifetime was observed from 50 to 60 °C, reflecting the thermal unfolding of STARD1 and the concomitant ligand dissociation (Fig. 6D,E). The half-transition temperature (~54 °C) was consistent with the results obtained by steady-state fluorescence measurements (Fig. 5A). At the same time, there was no significant changes in the lifetimes of **3NC** in the region of half-transition temperature of STARD1 (Fig. 6D), and only a slight increase of slow component of fluorescence decay was observed at temperatures above 55 °C, when the protein is unfolded, and aggregation may occur (providing additional hydrophobic environment for nonspecific binding of 3NC). This is in favor of our hypothesis that **3NC** binds to STARD1 with lower affinity and the fluorescent group exposed to the solvent, and that this binding mode could not stabilize the protein against the heat-induced unfolding.

Possible **20NP** and **3NC** binding modes to native STARD1 are shown in Fig. 6B. These models, predicted by FlexAID docking and consistent with our experimental data, suggest the “C3 OUT” as the more preferential binding mode for these fluorescence analogues (Fig. 6B), however, we cannot exclude the ability of **3NC** to bind outside the cavity.

## 3. Conclusions

To the best of our knowledge, the present study describes the first detailed hydrodynamic analysis of STARD1 and its solution structure using the stable STARD1^S195E^ mutant preparation which was obtained thanks to the recently described original protocol. The results obtained here by small-angle X-ray scattering (SAXS) with in-line SEC separation, suggests that the monomeric STARD1, preserved in a non-aggregated state in a wide range of protein concentrations (0.3-10 mg/ml), demonstrates rather rigid/compact structure with limited flexibility, unlikely undergoing significant structural rearrangements postulated previously in order to describe its cholesterol-binding mechanism. In contrast to the less probable spontaneous unfolding of the C-terminal α-helix [17, 18, 28], contradicting our SAXS data, we could validate that movements of the Ω1 loop [31, 41] could be the only structural rearrangements necessary for cholesterol binding by STARD1 and its homologs. The solution structure of STARD1 appears similar to the recently obtained solution NMR structure of STARD6 [41].

Probing the cholesterol binding cavity of STARD1 using the series of NBD-labeled cholesterol analogues with different position of the fluorescent group and comprehensive set of biochemical and biophysical techniques allowed us i) to expand the repertoire of STARD1 ligands and understanding of selectivity of this protein, ii) to find the optimal NBD-ligand for other STARD1/cholesterol interaction studies, i.e., **20NP**, which gives much better fluorescence response than commercially available and commonly used **22NC** and **25NC**, iii) to validate its binding affinity and stoichiometry (1:1), and iv) to specify the most probable orientation of the cholesterol core in the cavity. The applicability of powerful time-resolved fluorescence lifetime spectroscopy approaches to study STARD1/ligand interactions and thermal stability of the corresponding complexes is also demonstrated. Apart from fundamental importance, all this information will be helpful for further studies to screen novel potential STARD1 ligands (both, occurring naturally or designed specifically) which could outcompete fluorescent **20NP** reporter, and for structural studies on STARD1/ligand complexes evading such an analysis so far.

## 4. Materials and methods

### 4.1. Materials

**22NC** was from Molecular Probes (Eugene, OR), **25NC** was from Avanti Polar Lipids (Birmingham, AL). Cholesterol, pregnenolone and cholestenone were obtained from Sigma-Aldrich (St. Louis, MO). NBD-labeled steroids were synthesized for this work as described [54], *via* steps of reductive amination of either pregnenolone or cholestenone and further reaction with 7-nitrobenzoxadiazole-4-yl chloride [55-57]. **20NP** was obtained as racemic mixture of two isomers, **20NPα** and **20NPβ**, which were separated as described in [38]. In this study only **20NPβ** (20*S*-; retention time of 30.0 min [38]) was used because it provided the most reproducible results, lower background fluorescence, and higher fluorescence yield upon STARD1 binding.

All NBD-labeled cholesterol analogues were dissolved in 96% ethanol to get 200-300 μM stock solutions (concentration was measured in cuvettes (0.5 cm) by absorbance at 470 nm using extinction coefficient of 21,000 M^−1^ cm^−1^). All other chemicals were of the highest purity and quality available. All water solutions were prepared on the milliQ (18.3 MΩ/cm) water and filtered through the 0.22 μm filters before use.

### 4.2. Protein expression and purification

Cloning and purification of the wild type human StAR domain (residues 66-285) as the maltose-binding protein (MBP) fusion cleavable by 3C protease was described earlier [38]. Briefly, the protein encoded by H-MBP-3C-STARD1_66-285_ plasmid was expressed in BL21(DE3) cells of *Escherichia coli* and purified by subtractive immobilized metal-affinity chromatography and size-exclusion chromatography. Due to the low intrinsic solubility of STARD1, it was important to use SEC as the final stage of purification which allowed separation of the individual STARD1 from a small fraction of aggregated species [38]. Alternatively, the protein was co-expressed in C41(DE3) cells with the catalytic subunit of mouse cAMP-dependent protein kinase protein kinase A (PKA) cloned into a compatible pACYCduet-1 vector [58]. Purification of the co-expressed and presumably phosphorylated STARD1 was performed using the same protocol.

Ser195 of STARD1, known to be the key residue phosphorylated by PKA [10], was replaced by either Ala or Glu to imitate unphosphorylated and phosphorylated state of this residue, respectively. To this end, 5′-CCGAGGC<u>GCC</u>ACCTGTG-3’ (S195A) or 5’- CCGAGGC<u>GAAA</u>CCTGTGTG-3’ (S195E) forward mutagenic primers (mutated codons are underlined) and the plasmid encoding the wild type protein were used. The integrity and correctness of the obtained constructs were verified by DNA sequencing. The mutants were expressed and purified according to the procedure developed for the wild-type STARD166-285 [38]. After SEC, proteins were almost homogenous according to SDS-PAGE [59], were flash frozen in liquid nitrogen and stored at −80 °C. Protein concentration was determined using Bradford microassay [60] and calibration curve build by BSA standard solutions (ThermoFischer Scientific Inc.).

### 4.3. Native gel-electrophoresis in the presence of urea

In order to reveal charge differences in case of STARD1 and its mutants irrespective the shape factor, we used native gel-electrophoresis at pH 8.6 in the presence of 8 M urea in gels and in loading buffer. The samples (1 mg/ml) were incubated in the loading buffer and then subjected to electrophoresis according to the described procedure [61, 62].

### 3.4. Circular dichroism spectroscopy

CD spectra of STARD1^S195A^ and STARD1^S195E^ (1 mg/ml) were recorded at 20 °C on a Chirascan circular dichroism spectrophotometer (Applied Photophysics) as described earlier [38].

### 4.5. Steady-state fluorescence spectra measurements

All steady-state fluorescence spectra were recorded on a Cary Eclipse fluorescence spectrophotometer (Varian Inc.) equipped with a temperature controller. Protein samples (0.1-2 μM) were prepared on a buffer F1 (20 mM Tris-HCl (pH 7.5), 200 mM NaCl, 0.1 mM EDTA) or buffer F2 with Tris replaced by Hepes. Intrinsic tryptophan fluorescence spectra of STARD1 species in the absence or in the presence of different NBD-labeled cholesterol analogues were recorded in the range of 305-575 nm upon excitation at 297 nm, whereas NBD fluorescence spectra were recorded in the range of 475-650 nm upon excitation at 460 nm. To monitor energy transfer from tryptophan to NBD, spectra were recorded in the full range 305-575 nm upon excitation at 297 nm. The slits width was 5 nm and typically absorbance at the excitation wavelength was less than 0.1 to exclude the effect of inner filter. All spectra presented were averaged (3-5 measurements) and buffer corrected.

### 4.6. Cholesterol-binding activity

STARD1^S195E^ samples (1 μM) in buffer F1 were pre-incubated at 37 °C, and then the intensity of either Trp or NBD fluorescence was recorded before or after each addition of small 5-1 μl aliquots of a ligand stock solution in ethanol, so that the final ligand concentration varied in a range of 0-4 μM and that of ethanol was <1 %. In some cases, the protein was titrated by one ligand (**20NP** or **3NC**) and, after saturation was reached, was titrated by another one (**20NP** or cholesterol) to reveal competition for the binding site. After each addition, samples were mixed by microsyringe and equilibrated for 5 min at 37 °C prior to measurements. We did not observe substantial differences between 5 and 30 min pre-incubation. Alternatively, the series of probes with varying STARD1/ligand ratio was pre-incubated overnight in the fridge for full equilibration, but this gave almost the same results as the first approach. The experiment was repeated at least three times for each ligand, and the most representative results are shown and discussed.

To assess apparent binding parameters, the binding curves against total added ligand concentration (x) were fitted using a quadratic equation as described in [38, 63]:

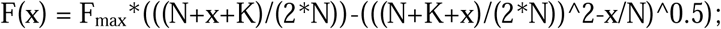

where F_max_ is fluorescence intensity (a.u.) at saturation, N - concentration of binding sites (μM), KD – apparent dissociation constant (μM), and F – fluorescence intensity (a.u.) at x ligand concentration.

Calculations and non-linear curve fitting were performed using Origin 9.0 Pro software.

### 4.7. Analytical size-exclusion chromatography (SEC)

SEC was used to analyze the oligomeric state of STARD1 at different concentrations and to validate physical interaction of STARD1 with the NBD-ligands. In the first case, STARD1^S195E^ samples at protein concentration in the range 0.3-2.8 mg/ml were pre-incubated for 30 min at room temperature and then loaded on a Superdex 200 Increase 10/300 column (GE Healthcare) equilibrated with buffer F1 and operated at 1.5 ml/min. In the second case, STARD1 (10 μM) was pre-incubated for 30 min at 37 °C in the presence of either **20NP**, **22NC**, **25NC**, or **3NC** (12 μM), and then subjected to SEC under the same conditions. In this experiment, dual wavelength detection (280 and 460 nm) was used to obtain protein-specific and ligand-specific profiles.

### 4.8. Small-angle X-ray scattering data collection and processing

The STARD1^S195E^ mutant was analyzed by synchrotron SAXS at the P12 beamline (PETRA III, DESY Hamburg, Germany) using an inline HPLC system for sample separation immediately preceding data collection. The sample (10 mg/ml; 75 μl) with no signs of protein precipitation was loaded on a Superdex 200 Increase 10/300 column (GE Healthcare) preequilibrated with filtered and degassed 20 mM Tris-HCl buffer (pH 7.6) containing 150 mM NaCl, 0.1 mM EDTA, 2 mM dithiothreitol, and 3 % glycerol. Separation was achieved at a 0.5 ml/min flow rate, and the flow was equally divided between the SAXS and TDA detection modules to ensure simultaneous data collection from the equivalent parts of the profile. TDA allowed simultaneous analysis of the eluate by absorption at 280 nm, refractive index (RI), and right angle light scattering (RALS). The RALS data for STARD1 and those for BSA standard were used to obtain Mw distribution for STARD1 over the profile (dn/dc was taken as 0.185). SAXS data frames (exposure time – 1 s, dead time – 1 s, wavelength – 1.24 Å) for the buffer (frames 500-1500) and the sample (frames 1880-1940) were collected. The time course revealed no radiation damage for protein frames. All buffer frames were averaged and subtracted from each protein frame. Protein frames were then scaled to the curve corresponding to the peak maximum and averaged by PRIMUS [64] to produce the resulting SAXS curve to be utilized for overall hydrodynamic parameters (I_0_, R_g_, D_max_, Porod Volume) assessment and modeling of the STARD1 solution structure using ATSAS package (http://www.embl-hamburg.de/biosaxs/software.html). Volume-of-correlation, V_c_, was determined as the ratio of the zero angle scattering intensity I_0_ to the total scattered intensity defined as an integral of S × I(S) versus S, up to S = 0.3 Å^−1^. The obtained V_c_ value was used to calculate M_W_ using an empirical relationship: M_W_ = Q_R_ / 0.1231, where Q_R_ = V_c_^2^ × R_g_^−^ [42].

*Ab initio* molecular envelope was built by averaging 20 independent DAMMIF [65] models (S ≤ 0.3 Å^−1^) using DAMAVER [66]. As the first approximation in structural model building, we used the only crystal structure of STARD1 available (PDB entry 3P0L chain A), and modified it by adding the missing parts (loop Ω1 residues, C-terminal residues 277-284, and the vector-derived N-terminal residues GPGS) and mutating Ser195 by phosphomimicking Glu residue using Coot [45] and i-Tasser [67]. Theoretical SAXS curves and fitting to the experimental data were calculated using Crysol [68]. The structural flexibility of STARD1 was analyzed using ensemble optimization method (EOM) [69]. Principles of building alternative models are described in the text.

HydroPro 1.0 (http://leonardo.inf.um,es/macromol/programs/hvdropro/hvdropro.htm) was used to independently estimate hydrodynamic parameters of STARD1 based on its crystal structure (PDB entry 3P0L, chain A, with our modifications described above). To this end, partial specific volume 0.73 cm^3^/g, solution density 1 g/cm^3^, Mw 24,800 Da, shell-model from atomic level (radius of elements 2.84 Å), were used for calculations.

### 4.9. Time-resolved fluorescence measurements

Fluorescence decay kinetics were recoded using time-and wavelength-correlated single photon counting setup based on SPC-150 module and HMP-100-50 detector (Becker&Hickl, Germany). Samples were excited using 450 nm laser diode (InTop, Russia) delivering 30 ps (FWHM) pulses, driven at a 10 MHz repetition rate or in CW mode. Necessary spectral band with a maximum near 540 nm was selected by ML44 monochromator (Solar, Belarus).

Fluorescence decay curves were approximated by a sum of exponential decay functions with the SPCImage (Becker and Hickl, Germany) software package. To compare different decay curves, we calculated the average decay time according to the following expression:

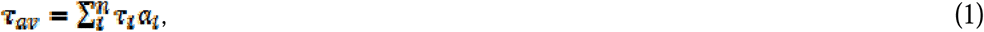

where τ _*i*_ and *a_i_* are the lifetime and the amplitude (normalized to unity: **Σ*_i_^n^ a_i_* = 1**) of the i-th fluorescence decay component, respectively.

During the experiment temperature of the sample was controlled by a cuvette holder Qpod 2e (Quantum Northwest, USA).

### 4.10. Docking simulations

Docking simulations of cholesterol and its fluorescent analogues into human STARD1 structure (PDB entry 3P0L, chain A with modifications; see above) were performed either in NRGsuite plugin for PyMol for run real-time docking with FlexAID [53] or using Autodock 4.2 with AutoDockTools [52].

The first approach is based on shape complementarity between ligands and receptors, accounts for protein side chain and ligand flexibility, but does not take into account electrostatics. Assignment of the binding site was done using GetCleft tool of the NRGsuite (minimum probe radius – 1.6 Å, hydrogens added, volume of the binding cavity to accommodate a ligand anchor atom estimated as 450 Å^3^), and the residues from the interior of the cholesterol binding cavity were treated as flexible. The simulations run used the genetic algorithm (1000 chromosomes, 1000 generations, share fitness model and the rest default parameters).

For the second approach, the protein structure was processed, and Gasteiger partial charges [70] were calculated and assigned to the atoms. The docking space was defined as a 60×60×60 Å^3^ box close to geometric center of the protein. Ligands were created and prepared using HyperChem 7.01 (Hypercube). The Lamarckian genetic algorithm with default parameters was applied for rigid docking calculations. The binding energy values were calculated automatically by Autodock.

## 5. Acknowledgements

Authors are thankful to Dr. Alexey Mantsyzov (Moscow State University) for help in calculation maximal RMSD in the STARD6 ensemble and to Dr. Cy Jeffries (EMBL Hamburg) for assistance during SAXS data collection and initial data processing. This investigation was supported by the Russian Foundation for Basic Science (grant 17-04-00331a to N.N.S.) and the Program “Molecular and Cell Biology” of the Russian Academy of Sciences (N.N.S.). E.M.G. acknowledges the funding by Russian Foundation for Basic Science and Moscow city Government according to the research project № 15-34-70007 «mol_a_mos». Ya.V.F. acknowledges the grant from the Belarusian State Program of Scientific Investigations (№ 20161380).

---

**Figure S1.**
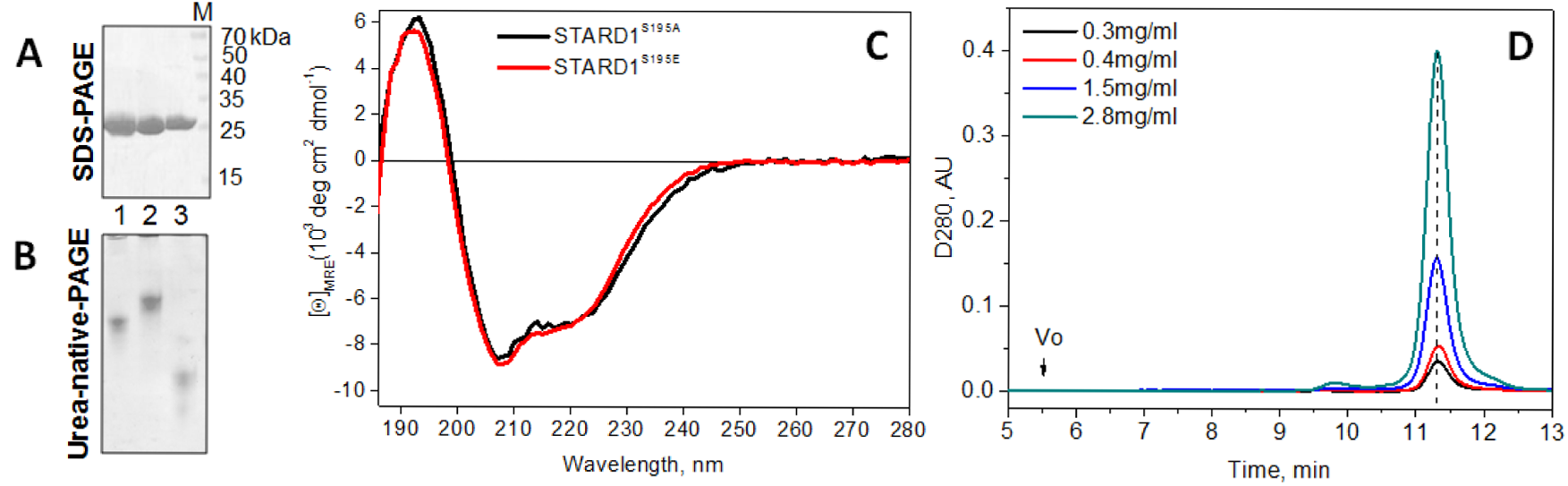
Properties of STARD1 preparations obtained according to the procedure based on the cleavable His-MBP tagged fusion. Analysis of electrophoretic purity and mobility of STARD1^S195E^ (1), STARD1^S195A^ (2), or wild-type STARD1 bacterially co-expressed with PKA (3) by SDS-PAGE (**A**) or Native-PAGE at pH 8.6 in the presence of urea (**B**). **C**. Far UV-CD spectra of STARD1^S195E^ and STARD1^S195A^ (1 mg/ml) at 20 °C. **D**. Analytical size-exclusion chromatography of STARD1^S195E^ at different protein concentrations in the loaded sample (indicated). Vo shows position of the void volume, where no STARD1 aggregates were found even at the highest concentration tested. The vertical dashed line connects the maxima of the peaks at different concentrations.

**Fig. S2.**
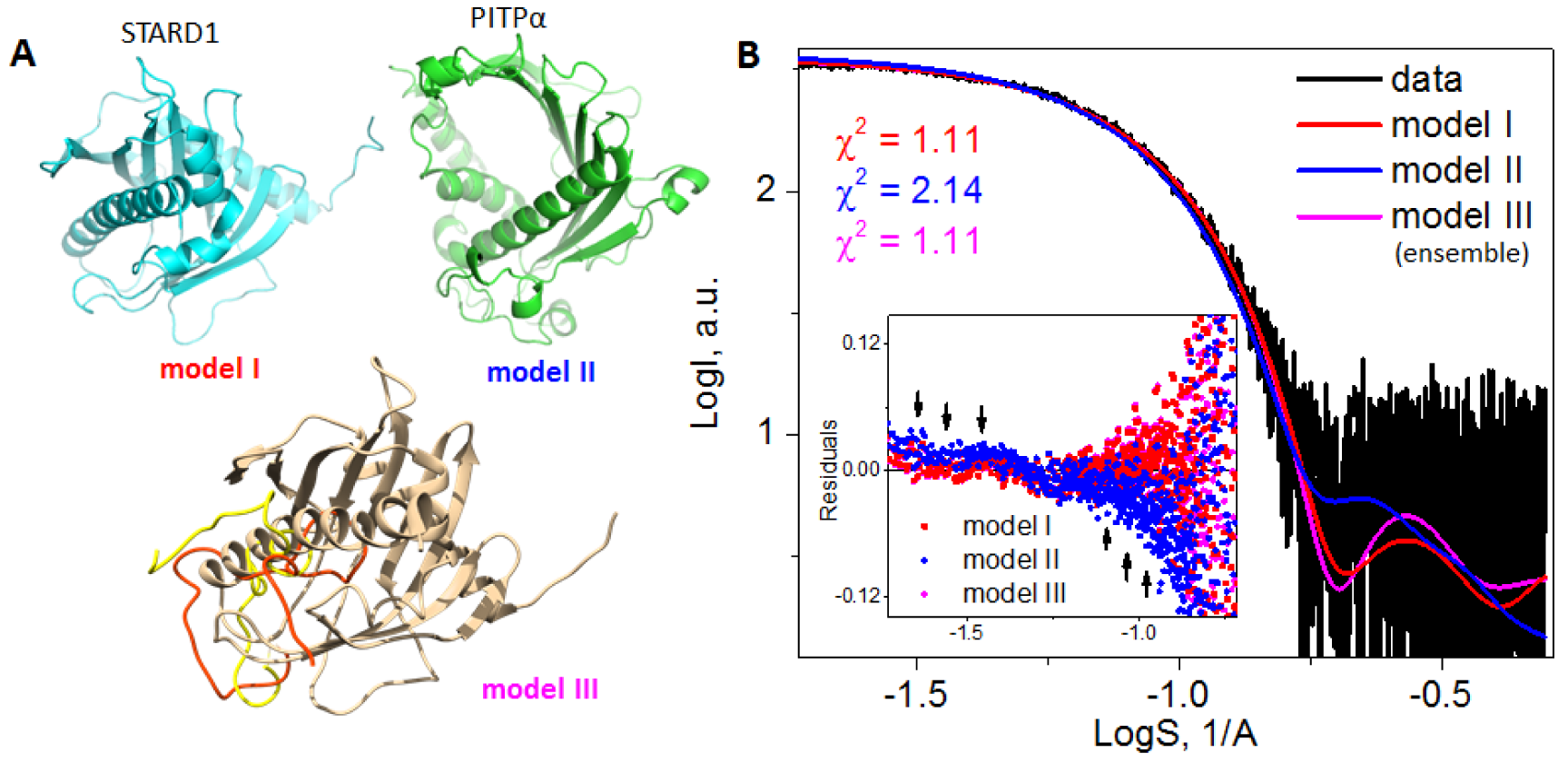
Alternative models of STARD1 in solution. **A.** Structural representation of the models I, II, and III. Model I corresponds to the modified crystal structure of STARD1 (see text), model II is the crystal structure of PITP_α_ protein (=STARD10) containing the START domain in an “open” conformation (PDB entry 1KCM), representing the “clam-shell” like opening as the gating mechanism to bind cholesterol. Model III corresponds to the best fitting ensemble of hypothetical structural models with the unfolded C-terminal α4-helix (residues 250-285), as proposed by [17, 18, 28]. The initial position of the α4-helix (beige) and two selected partially unfolded conformers (red, yellow) are shown. Note that out of 10,000 created models sampling all possible positions of the unfolded α4-helix, only the most compact ones with the unfolded segment lying within the cavity are selected. **B.** Fits by the models I-III to the experimental SAXS data. Inset: residuals at the low-angle region showing discrepancies in the case of model II (arrows indicate misfits). Note that the fit from model III was obtained by ensemble modeling. Drawn using PyMol 1.69 (models I and II) and Chimera 1.11 (model III).

**Fig. S3.**
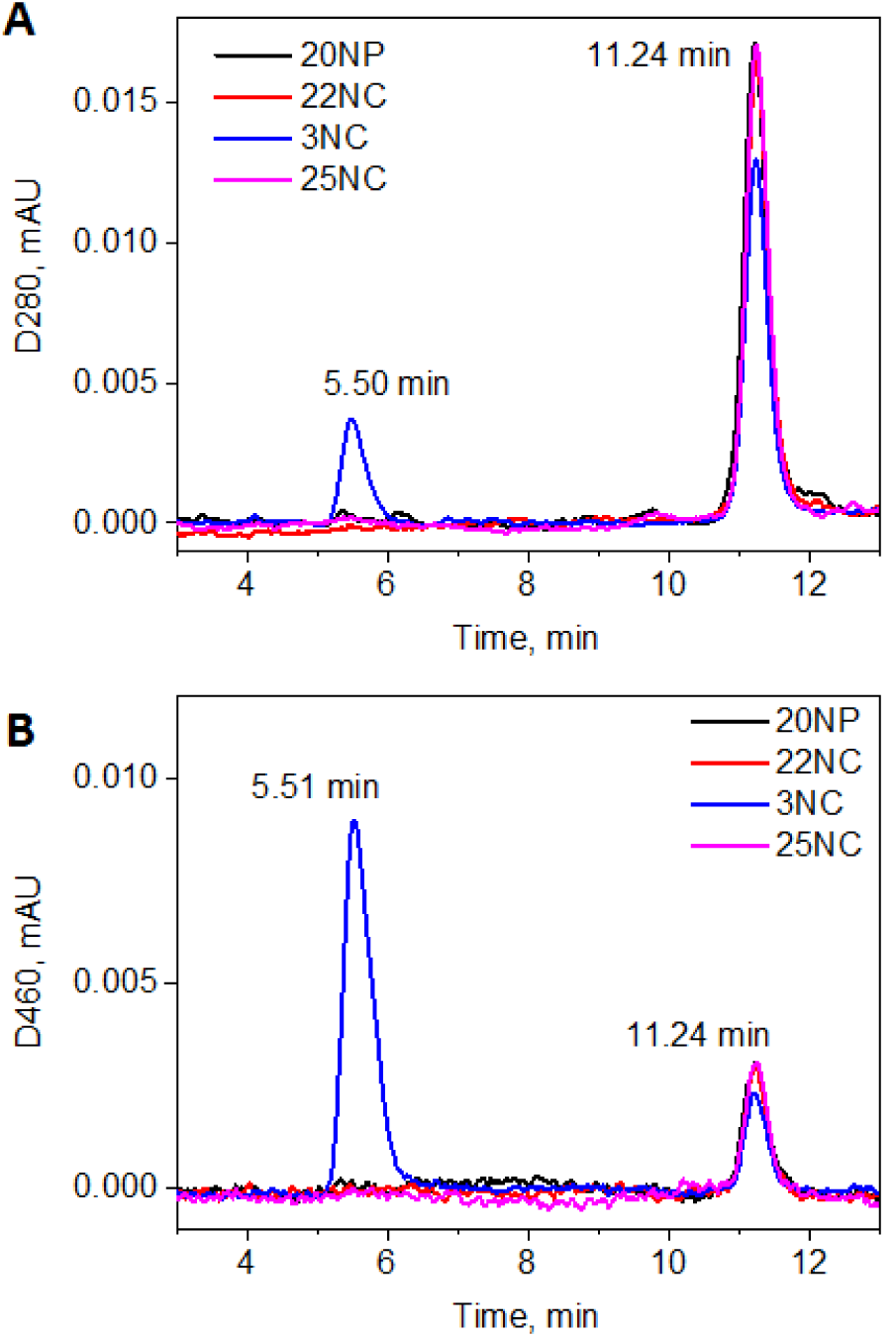
Validation of physical interaction between the NBD-ligands and STARD1^S195E^ by analytical size-exclusion chromatography on a Superdex 200 Increase 10/300 column followed by either protein-specific (**A:** 280 nm) or ligand-specific (**B:** 460 nm) absorbance. Elution times of the peaks corresponding to aggregated (~5.50 min) and monomeric (11.24 min) STARD1 are shown. Flow rate 1.5 ml/min. The experiment was repeated twice with substantially similar results.

**Fig. S4.**
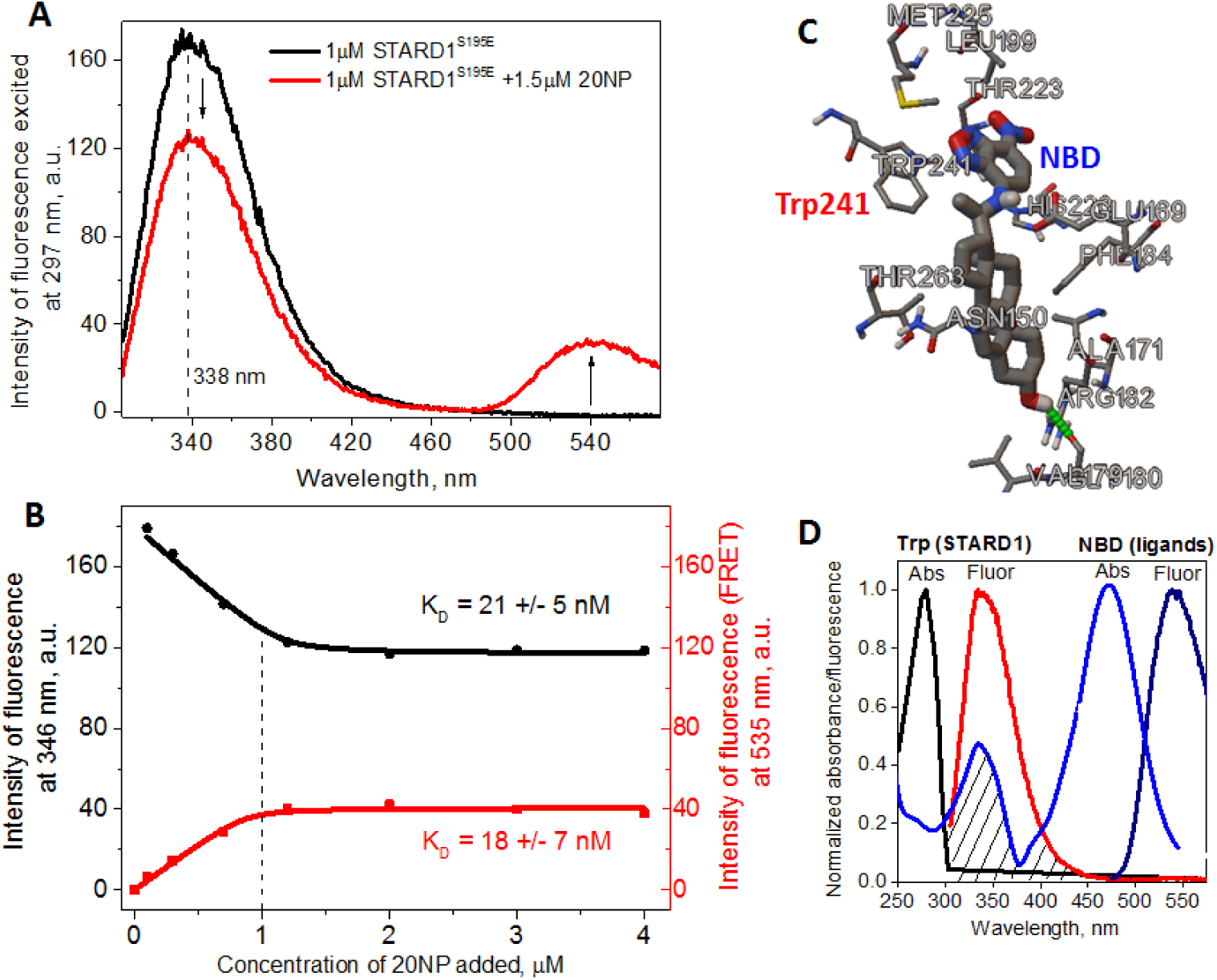
FRET from STARD1 tryptophans to the NBD group of bound 20NP. **A**. Spectral changes upon addition of 20NP (indicated by arrows). Excitation at 297 nm, slits width 5 nm, temperature 37 °C. Maximum of Trp fluorescence is indicated by dashed line. **B**. Titration of STARD1^S195E^ solution (1 μM) by increasing concentrations of 20NP followed by either quenching of tryptophan fluorescence (black) or increase in the intensity of NBD fluorescence (red). The vertical dashed line corresponds to the 1:1 ratio. **C**. Docking results in Autodock showing the proximity of Trp241 and the NBD-group of the bound 20NP and possibility of FRET (arrow). **D**. Spectral overlap in the fluorophore system involved.

**Fig. S5.**
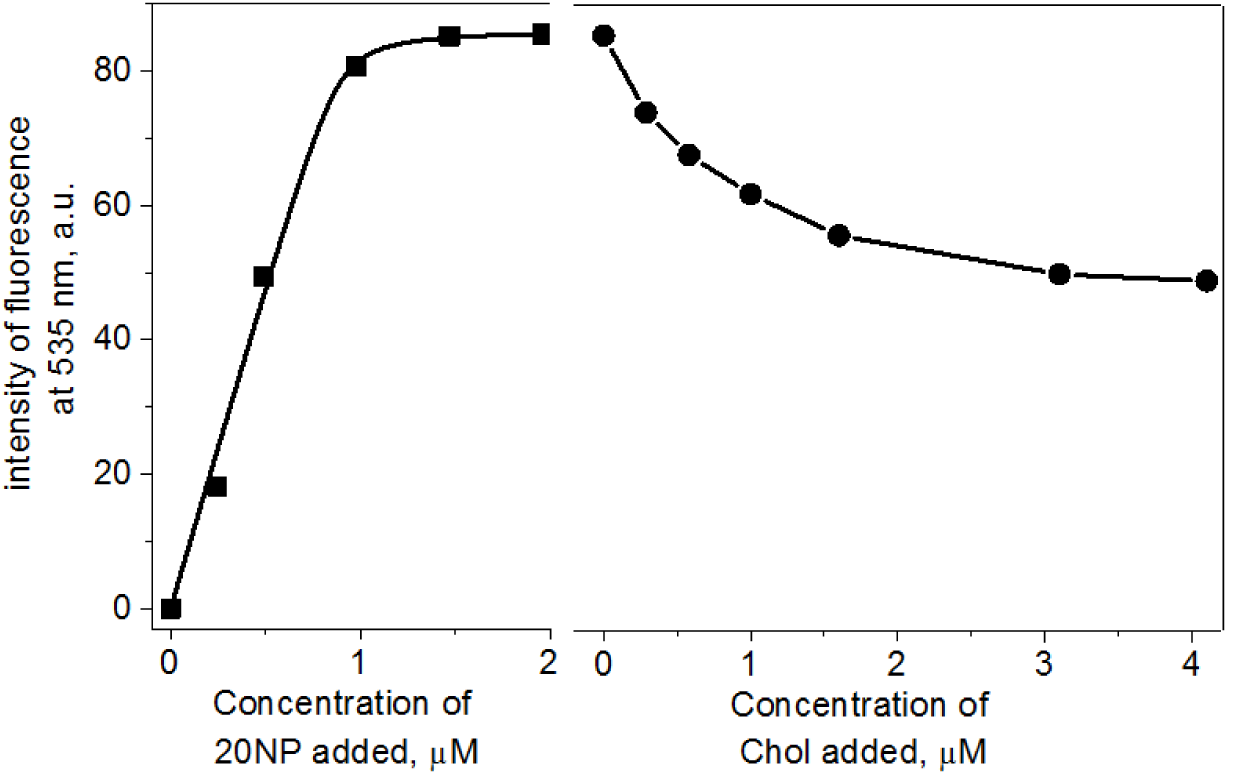
Competitive replacement of STARDl-bound 20NP by cholesterol. STARD1^S195E^ (1 μM) was first titrated by 20NP until saturation (2 μM) (squares) and then by increasing amounts of cholesterol (circles). Fluorescence was excited at 460 nm. Temperature was 37 °C. Note that concentrations of cholesterol were significally higher than its critical micelle concentration (tens nM), therefore approximation of the curve and assessment of binding parameters were not possible.

**Fig. S6.**
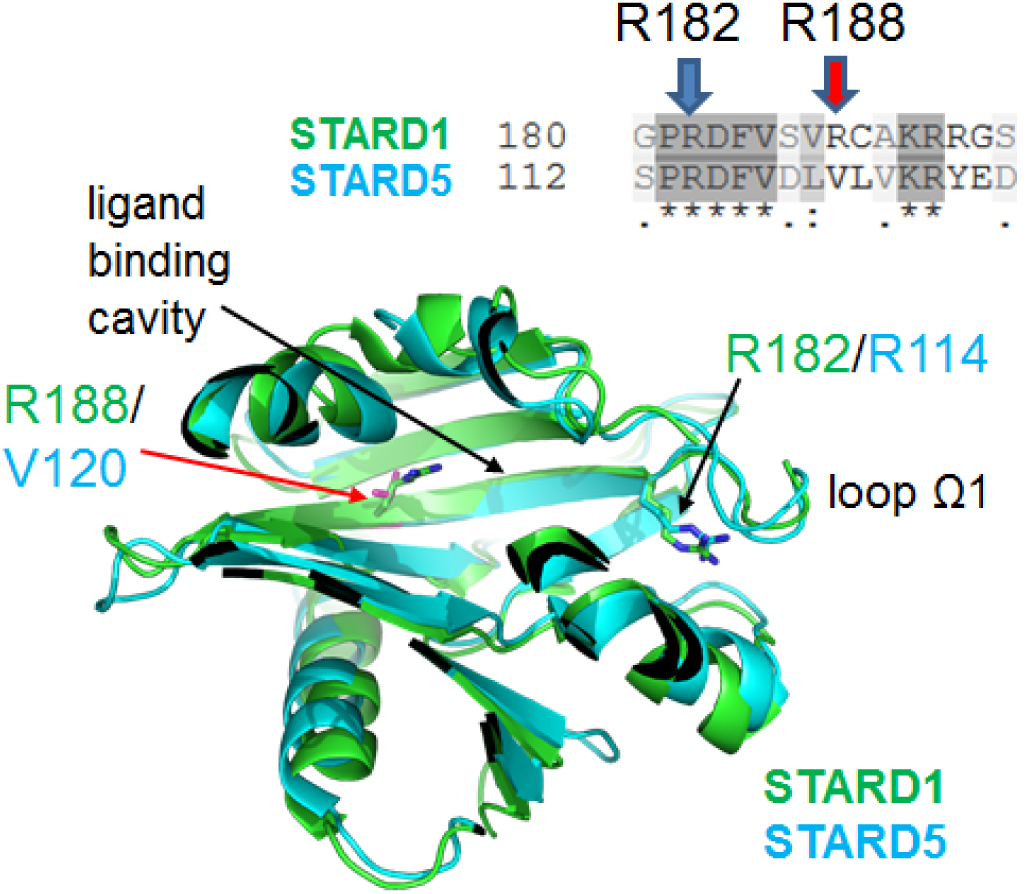
Alignment and superposition of STARD1 and STARD5 structures exemplifying lesser conservativity of the R188 position. The primary structure was aligned using Clustal Omega, the level of homology is shown as greyscale. The crystal structures of STARD1 (PDB entry 3P0L) and STARD5 (PDB entry 2R55) were superimposed and drawn in PyMol 1.69 (sagittal plane), with the main features highlighted.

**Fig. S7.**
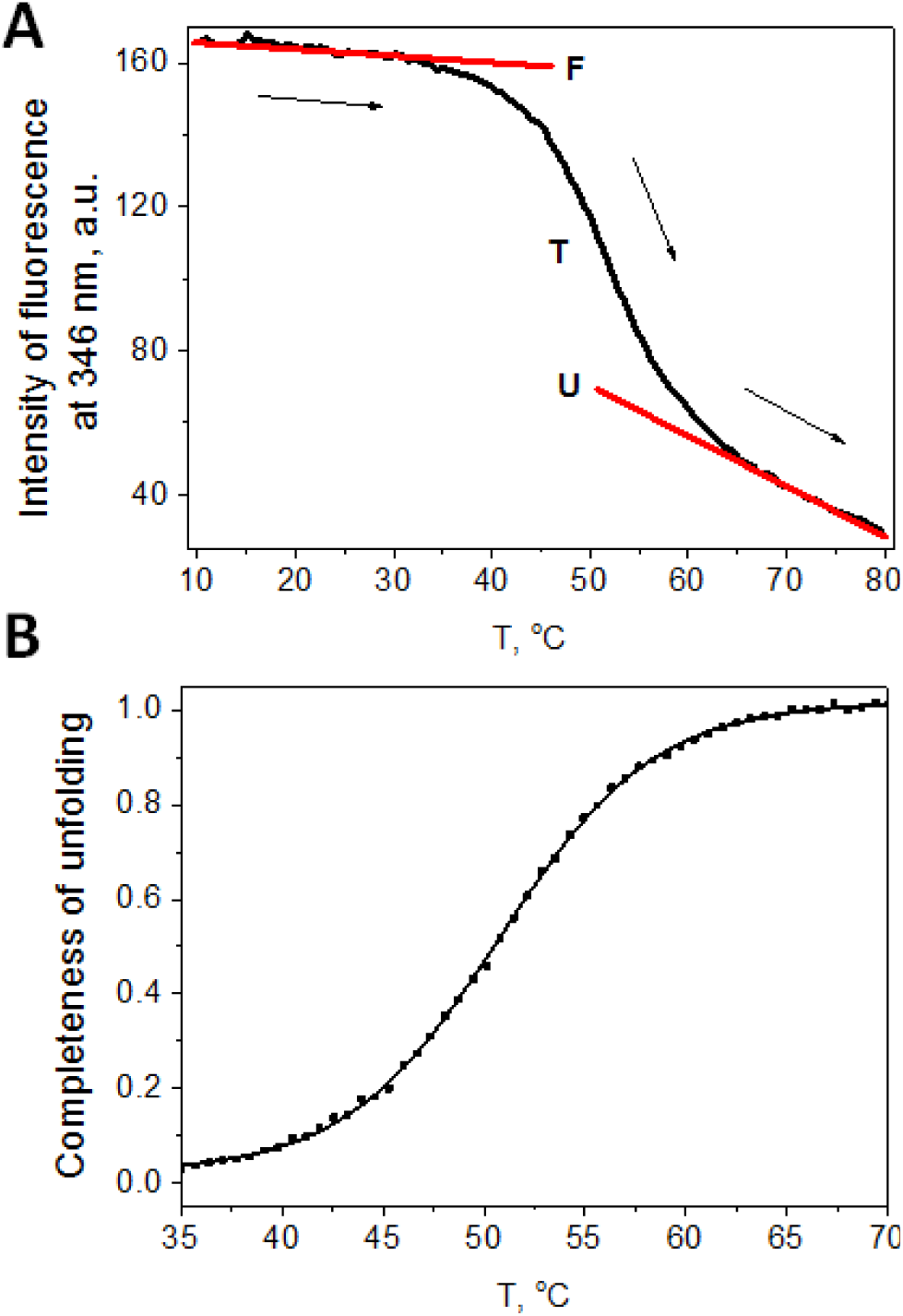
Thermally-induced changes in the intrinsic tryptophan fluorescence of STARD1^S195E^ The sample was heated at a constant rate of 1 °C/min and changes in the intensity of fluorescence excited at 297 nm were registered at 346 nm in a range of 10-80 °C. **A**. Raw data with the three regions corresponding to the (F)olded, (T)ransition, and the (U)nfolded state, highlighted. Arrows indicate the direction of heating. **B**. The same data converted to a form of temperature dependence of the so-called completeness of unfolding allowing to determine the corresponding half-transition temperature (T_0.5_).

